# Skin surface biomarkers are associated with future development of atopic dermatitis in children with family history of allergic disease

**DOI:** 10.1101/2023.07.11.548501

**Authors:** Takahiro Sato, Janet Nikolovski, Russell Gould, Imane Lboukili, Pierre-Francois Roux, Gabriel Al-Ghalith, Jeremy Orie, Richard Insel, Georgios N. Stamatas

## Abstract

**Background:** Atopic dermatitis (AD) is a common childhood chronic inflammatory skin disorder that can significantly impact quality of life and has been linked to the subsequent development of food allergy, asthma, and allergic rhinitis, an association known as the “atopic march.”

**Objective:** The aim of this study was to identify biomarkers collected non-invasively from the skin surface in order to predict AD before diagnosis across a broad age range of children.

**Methods:** Non-invasive skin surface measures and biomarkers were collected from 160 children (3-48 months of age) of three groups: (A) healthy with no family history of allergic disease, (B) healthy with family history of allergic disease, and (C) diagnosed AD.

**Results:** Eleven of 101 children in group B reported AD diagnosis in the subsequent 12 months following the measurements. The children who developed AD had increased skin immune markers before disease onset, compared to those who did not develop AD in the same group and to the control group. In those enrolled with AD, lesional skin was characterized by increased concentrations of certain immune markers and transepidermal water loss, and decreased skin surface hydration.

**Conclusions:** Defining risk susceptibility before onset of AD through non-invasive methods may help identify children who may benefit from early preventative interventions.

## INTRODUCTION

Atopic dermatitis (AD) is a common chronic inflammatory skin disorder that often manifests within the first year of life^1^ and can significantly impact the quality of life of the child and the caregiver.^2^ The incidence of AD in children ranges from 15 to 30%.^3^ AD with onset in early life has been linked to the subsequent development of food allergy, asthma, and allergic rhinitis, an association known as the “atopic march”.^4^ This progression is dependent on various underlying factors such as genetic mutations, skin barrier impairments, immune dysfunction, and environmental conditions.^5-8^

Children with a family history of allergic disease are at highest susceptibility for developing atopic diseases.^9,10^ Previous studies have identified different phenotypes of AD with different trajectories of disease progression and risk of development of atopic comorbidities.^11^ These phenotypes are characterized by the age of onset of the first symptoms (two with early onset [transient or persistent progression] and one late onset [after 2 years]) and the natural course of the disease from birth through early childhood. Within these unique phenotypes, there is emerging evidence of differential association with risk factors, but risk factors and prediction markers for late-onset AD remain unknown.

There is a need to define risk susceptibility biomarkers that can be used to identify children before onset of AD, who may benefit from preventative interventions. Importantly, capturing these biomarkers must be done using minimally invasive methods, since biopsies and blood draws are not always practical in infants/children. Local skin barrier biology and underlying mechanisms have emerged as an area with significant predictive potential insight.^12^ Tape stripping has shown promise in detecting cutaneous gene expression/protein biomarkers, and associations have been found between disease severity and Th2 and Th17/Th22 products in lesional and non-lesional AD skin.^13,14^ Moreover, microbiome signatures of the skin have been encouraging for diagnosing AD^15^ and were associated with AD severity^16^ and risk of developing AD.^17,18^

The aim of this study was to identify skin surface biomarkers of healthy individuals that are associated with future onset of AD. Such biomarkers could also help gain a deeper understanding of disease mechanism. Skin surface samples were collected from: (A) healthy children with no family history of allergic disease (healthy/low-risk), (B) healthy children with family history of allergic disease (healthy/high-risk), and (C) children with active AD lesions. Noninvasive data collected included measures related to skin function and skin composition.

## MATERIALS AND METHODS

### Study design

A single-center, exploratory study was conducted to compare skin biomarkers in 3 groups of children aged 3-48 months, based on their status of AD and their family history of atopic disease. The study was reviewed by an independent institutional review board for ethical approval (study 19.0457-39, certificate 2019/026, proDERM GmbH, Hamburg, Germany). All study participants were residents of the Hamburg, Germany area. The study was conducted according to the principles of the Declaration of Helsinki. All parents/legal authorized representatives (LARs) of the participating children had a complete understanding of the test procedure and gave written consent to the study. Prior to the study, the parents/LARs were instructed not to apply any leave-on cosmetics on their child in the test area within the last 3 days prior to the start of the study or apply any cleansing products on their child in the test area on the day of the measurements. They were instructed to avoid contact of the test area with water within the last 2 hours prior to instrumental measurements.

Study participants were divided into the following groups: (A) 29 healthy/low-risk, defined as without history of AD in self or family (parent or siblings), hay fever, or allergic asthma; (B) 101 healthy/high-risk, defined as without AD but with family (parent or siblings) history of allergic disease (AD, hay fever or allergic asthma); (C) 30 with active AD with local SCORAD index of 4-8 as assessed by a study pediatrician on the atopic lesion area by rating 7 parameters (erythema, edema and papules, weeping and crusts, excoriation, lichenification, dryness, and pruritus), according to the following scale: 0 = None, 1 = Slight, 2 = Moderate, 3 = Strong. The subjective parameter pruritus was assessed by the LARs of the participating children. Note that Group B was intentionally oversampled (twice as that of either groups A or C) since we could not accurately predict up front how many children of that group would convert to AD within 12 months.

Children with serious illnesses or systemic medication past or present, any skin related conditions, including AD (besides for group C), seborrheic dermatitis, psoriasis or skin infections were excluded from the study.

At 6 months after their visit, parents/LARs were contacted to fill out a Health History Questionnaire. At 12 months, parents/LARs of participants in groups A and B were contacted by phone call to inquire if their child was diagnosed with AD since the first visit.

### Measurements

All measurements took place in an air-conditioned room at a temperature of 21±1°C and at 50±5% relative humidity. Before measurements, study participants stayed in the climatized room for ≥30 min.

Skin surface hydration (SSH) was evaluated using the electrical capacitance method (Corneometer CM825, Courage & Khazaka, Cologne, Germany), with 5 measurements at each test area.

Skin barrier was assessed by measuring the rate of transepidermal water loss (TEWL) with a closed chamber system equipped with a condenser (Aquaflux AF200, Biox Systems Ltd., London, UK) with one measurement at each test area. This instrument guarantees the recording of a stabilized value, an aspect that removes the need for multiple measurements.

Photos of the test area were acquired using a high-resolution digital camera (Canon EOS 5D Mark III; 50 mm Macrolens/28 mm Macrolens) in combination with a dermatoscope (DERMlite TM Foto, 3Gen Inc., San Juan Capistrano, CA, USA) with ring-light illumination and distance holder. The captured test area was 1.5-2 cm/2-2.5 cm (28 mm lens) in diameter. White balancing and color calibration were performed using the X-Rite Color Checker.

Raman spectra at consecutive depths in the skin were acquired using an *in vivo* confocal Raman microspectrometer (gen2-SCA Skin Analyzer, River D, Rotterdam, The Netherlands). Concentration profiles of water (wavenumber range 2600-3800 cm^-1^) and NMF components (wavenumber range 400-1800 cm^-1^) were calculated from the Raman spectra using manufacturer’s software (SkinTools). For water profiles, spectra were acquired in the high wavenumber spectral region from the skin surface up to a depth of ∼48 μm in the skin, in steps of 2.0 μm with an integration time of 1 sec and a 25 μm diameter pinhole, with ∼6 profiles per test area. For amino acid profiles, spectra were acquired in the fingerprint spectral region from the skin surface up to a depth of ∼48 μm in the skin, in steps of 2.5 μm with an integration time of 5 sec and a 100 μm diameter pinhole, with ∼6 profiles per test area. Depth profiles for the resistance to water transport were calculated as previously described.^19^

Immune marker analysis was performed by swabbing the skin areas of interest using specialized swabs soaked in buffer (FibroTx, Tallinn, Estonia). Samples were stored on dry ice until further processing and at -80°C until shipment to FibroTx for marker analysis using a spot enzyme-linked immunosorbent assay. Samples were analyzed for the presence of chemokine (CC) ligand (CCL)-17, CCL-2, CCL-22, CCL-27, CXC chemokine ligand (CXCL)-1/2, human *β*-defensin (hBD)-1, interleukin (IL)-18, IL-1α, IL-1*β*, IL-1 receptor antagonist (IL-1RA), IL-36*γ*, IL-8, S100A8/9, and vascular endothelial growth factor (VEGF)-A. These chemokines were selected for their reported involvement in skin inflammatory processes and specifically in AD.^13-18^

Measurements by *in vivo* Raman confocal microspectroscopy were performed on the volar forearm (left or right side, randomized), and TEWL rates, SSH, imaging, and immune marker sampling were performed on the arm and center of the facial cheek (left or right side randomized). For group C, lesion assessments were preferably performed on the AD lesion and additionally on healthy skin beside the lesional skin (Suppl. Table S1). Skin swabs were preferably performed on the opposite side of the body (if not involved in AD) or next to the areas of the instrumental measurements.

### Statistical analysis

All statistical analyses were performed on R using the packages *DescTools, rstatix*, and *ggplot2*. Details of the statistical analyses are presented in the Supplemental Methods.

## RESULTS

### Participant characteristics

Children (N=160), aged 3-48 months, were enrolled in the study and divided into three groups: (A) healthy/low-risk group, (B) healthy/high-risk group, and (C) children with active AD lesions. All were Caucasian with a Fitzpatrick type between I-III. Baseline characteristics are shown in Table 1.

### Children in the high-risk group who developed AD within 12 months display differentiated expression of skin surface immune markers

In group B, 11 children presented with confirmed AD diagnosis within 12 months after skin assessment (converters), 71 reported no AD diagnosis (non-converters), and 19 were not reached. Of the 11 converters, 2 developed AD before 12 months of age (early-onset phenotype), 5 between 12–24 months (uncertain-onset phenotype), and 4 after 24 months of age (late-onset phenotype). There was no statistical difference between converters and non-converters in mode of birth, sex, age or in measured parameters of skin barrier function and SSH (Table 1, Suppl. Table S2).

A minimum of four immune markers with surface concentrations measured on the arm were required for a combined Z-score that was significantly different (p<0.05) between converters and non-converters (Fig. 1, Suppl. Table S3). The 4 selected markers for the combined Z-score (hBD-1, IL-1RA, IL-36*γ*, S100A8/9) were the ones ranking with lowest p-value on the arm data (Suppl. Table S2). The odds ratios (OR) of the combined Z-score for developing AD within 12 months were 19.2 for the arm and 9.1 for the face (Fig. 2, Suppl. Table S4).

**Figure 1:**
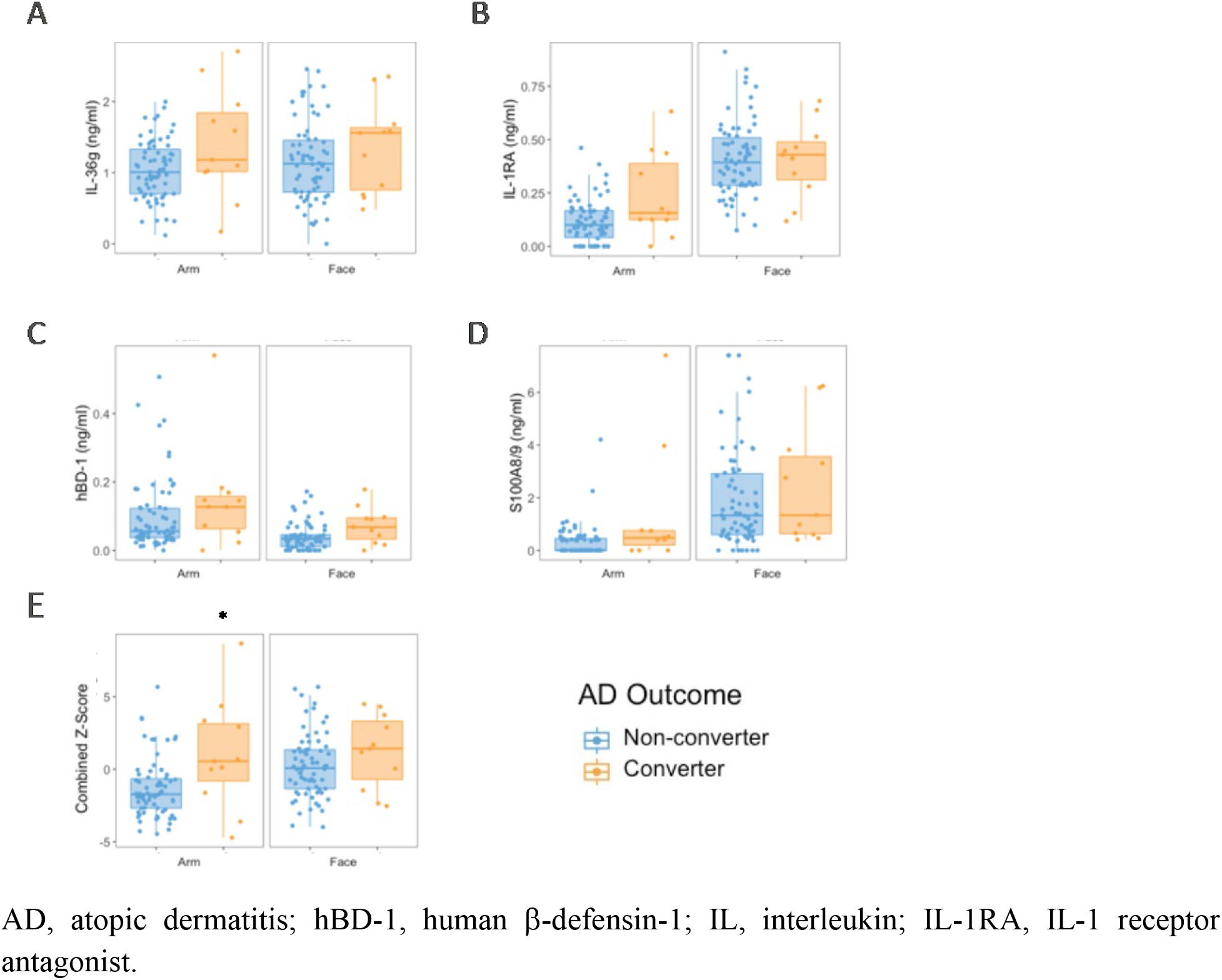
Skin surface immune marker concentrations measured on healthy children with family history of allergic disease who developed AD within 12 months (converters), compared to those who did not (non-converters). The surface concentrations of four markers associated with AD conversion are shown: (A) IL-36*γ*, (B) IL-1RA, (C) hBD-1, and (D) S100A8A/9. (E) Z-Scores of the surface concentration values of the 4 markers combined for each subject were statistically different between converters and non-converters for the arm site but not for the face (p<0.05).

**Figure 2:**
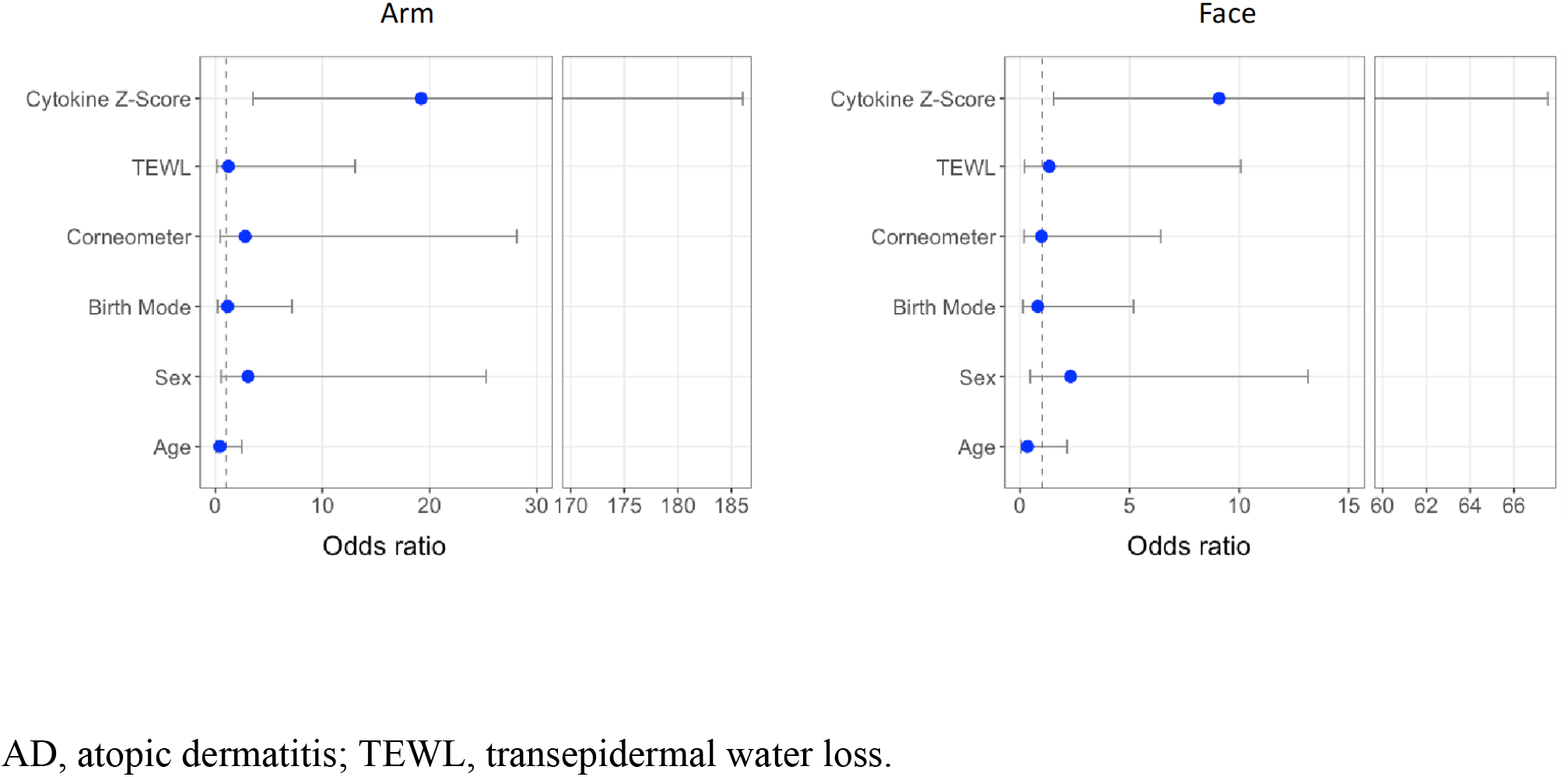
Odds ratios for developing AD within 12 months following skin assessment in healthy children with family history of allergic disease. Continuous variables were converted into binary high/low categories by classifying whether subjects were above the third quartile of that variable in healthy/low-risk subjects. Odds ratios were generated using logistic regression analysis.

Four different machine learning models trained on concentration data of all analyzed immune markers taken from skin surface swabs of AD participants (lesion) vs. healthy/low-risk participants showed high accuracy in separating the two groups as shown in receiver operating characteristic (ROC) curves. The median area-under-the-curve (AUC) values ranged from 0.93-1.00 for the arm and 0.75-0.94 for the face (Fig. 3A,B). These models were then tested on predicting healthy/high-risk converters vs non-converters and could predict development of AD on the arm with AUC values ranging from 0.65-0.78 for the arm (Fig. 3C). Immune marker data collected from the face had lower predictive ability as evidenced by AUC values ranging from 0.45-0.63 (Fig. 3D).

**Figure 3:**
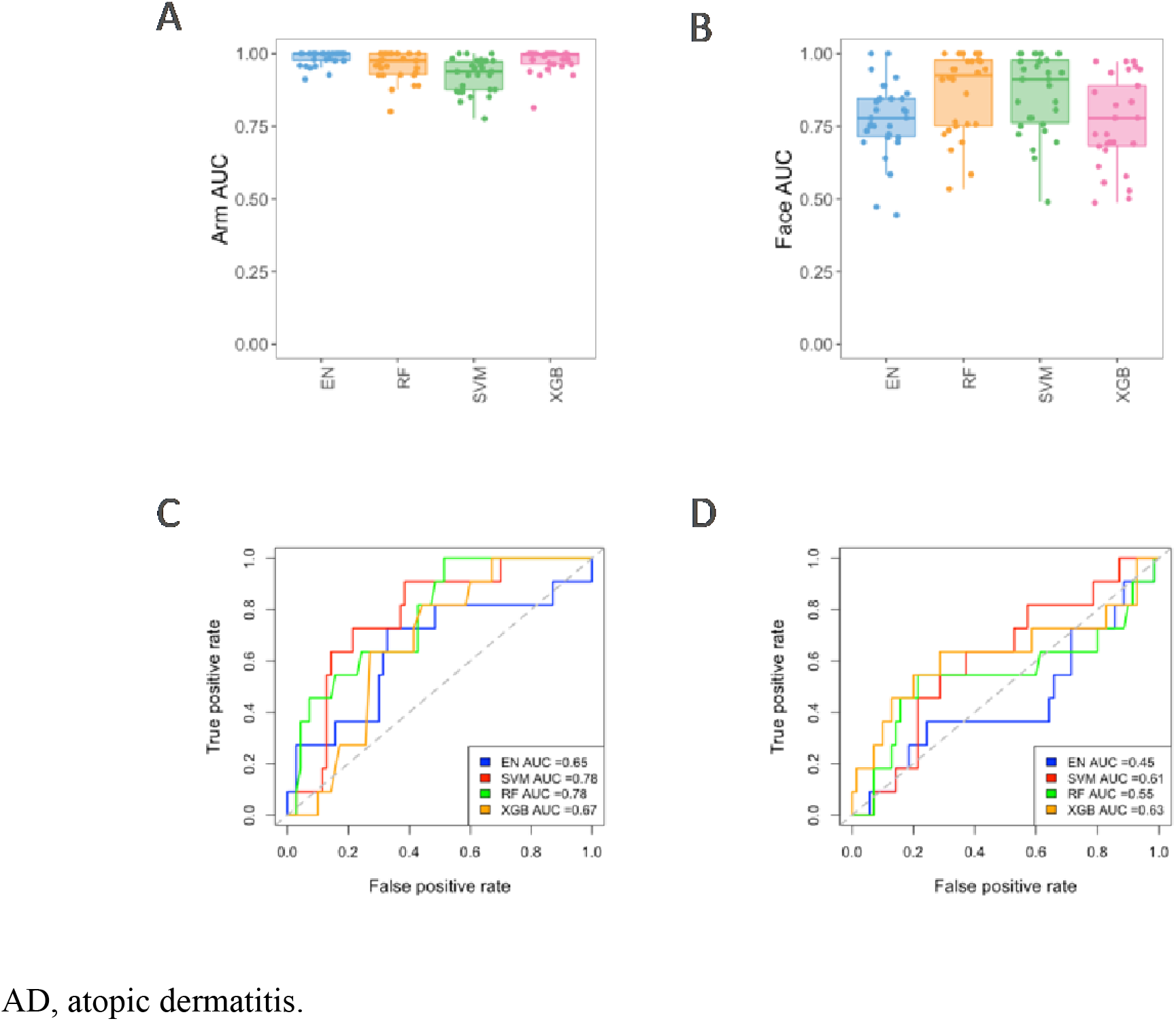
Receiver operating characteristic (ROC) curves for machine learning models predicting AD development within 12 months following skin assessment in healthy children with family history of allergic disease. Four different models: Elastic Net (EN), Support Vector Machine (SVM), Random Forest (RF), and XGBoost (XGB), were initially trained on AD lesional tissue vs healthy low risk using concentration data from all 14 markers and underwent 3-fold cross-validation repeated 10 times for hyperparameter tuning (A) on the arm and (B) on the face. These trained models were then tested on predicting converters vs. non-converters (C) on the arm and (D) the face.

Although the resistance to water transport was decreased for the healthy/high-risk and the AD group on non-lesional sites compared to the healthy/low-risk group, there was no statistical difference between converters and non-converters in the healthy/high-risk group (Fig. S1). Similarly, although the concentration of total amino acids in the *Stratum Corneum* (SC) was lower in the AD group on non-lesional sites compared to the healthy groups, there was no statistical difference between converters and non-converters in the healthy/high-risk group (Fig. S2).

### AD-lesional skin is characterized by increased expression of skin surface immune markers

Lesional AD sites displayed typical dryness and redness (Fig. S3) and impaired skin water barrier function on the arm site, as evidenced by significantly increased TEWL values and significantly lower SSH compared to healthy/low-risk participants (Suppl. Table S5). There was no statistical difference between the healthy/low-risk group and the AD group on non-lesional sites in TEWL or SSH values (Suppl. Table S5).

Immune markers measured on lesional AD skin had higher surface concentrations on both the face (7/14 markers upregulated) and the arm (6/14 markers upregulated) when compared to the skin of healthy/low-risk participants (S100A8/9 shown as example in Fig. 4A, Suppl. Table S5).

**Figure 4:**
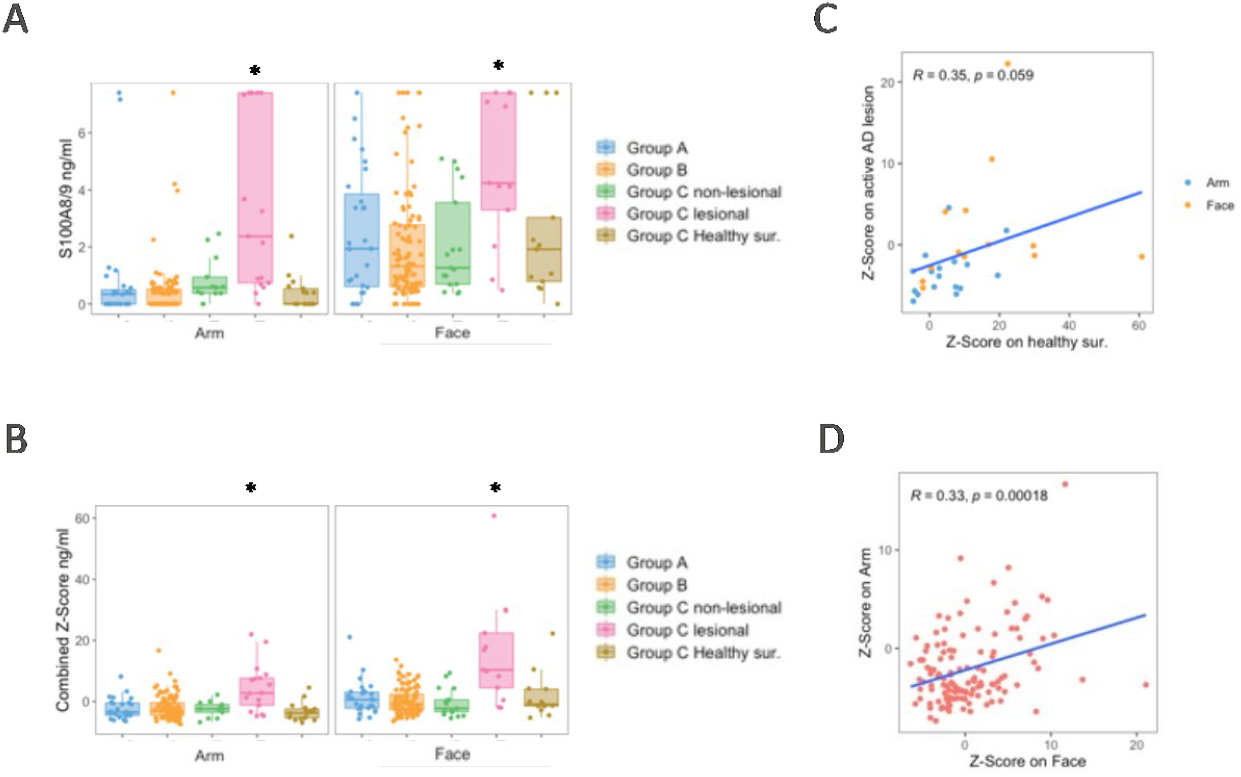
Comparison of skin surface concentration for immune markers between the studied groups. (A) The skin surface concentration data for S100A8/9 are shown as an example of a marker that had significant higher expression in AD lesional skin compared to healthy low-risk subjects. (B) Combined Z-score of the 14 immune markers had significant higher expression in AD lesional skin compared to healthy low-risk subjects. (C) Combined marker Z-scores in AD lesional tissue show a weak correlation with the corresponding values for healthy surrounding tissue for the same individual. (D) Combined marker Z-scores in healthy subjects on the arm show a weak but significant correlation with the corresponding values on the face for the same individual. Group A: Healthy/low-risk, Group B: Healthy/high-risk, and Group C: AD patients. * indicates statistical significance at p<0.05.

The combined Z-score (sum of Z-scores from 14 marker concentrations, a measure of general inflammation) was upregulated in AD lesions compared to healthy/low-risk participants on the face and arm (Fig. 4B). The combined Z-score was weakly correlated between active AD lesions and healthy surrounding tissue of these lesions, as well as between samples from the arm and face of healthy participants (Fig. 4C,D).

The concentrations of CCL-22 and CCL-27 positively correlated with TEWL on AD lesions on the arm, suggesting a potential role for these markers in being involved in the skin barrier dysfunction (Fig. S4).

## DISCUSSION

This study found that the combination of four skin surface immunity markers (IL-36*γ*, IL-1RA, hBD-1, and S100A8/9) is linked with increased risk of developing AD, particularly for the arm site. Moreover, the participants who developed AD exhibited different skin microbes at baseline compared with those who did not.

The interleukin cytokine family plays important roles in host defense, but excessive immune activation and altered receptor balance can lead to the development of chronic disease, such as AD. The biologic activity of pro-inflammatory IL-1 family agonists is mediated in part by the natural receptor antagonists IL-1RA, IL-36RA, IL-37, and IL-38.^20^ These four anti-inflammatory IL-1 family members are constitutively active and highly expressed in the epidermis by keratinocytes. The most important regulatory molecule for IL-1α activity is IL-1RA.^21^ Both IL-1RA and IL-36*γ* were found to be significantly upregulated in active lesions compared to normal controls in the skin of children with early-onset AD.^13^ Additionally, filaggrin loss-of-function mutations have been associated with enhanced expression of IL-1α cytokines in the SC of AD patients and in murine models, leading to increased IL-1RA.^20^ The observations here suggest that upregulation of IL-1RA and IL-36*γ* are early indicators for the onset of AD, relevant to the pathophysiology of the disease.

β-defensins, small cationic peptides secreted by keratinocytes, are important in cutaneous innate immunity.^22^ In particular, hBD-1 is expressed at relatively high levels following birth and gradually declines over the first three years of life,^23^ which was confirmed in this study (Fig. S5). The high levels of antimicrobial peptide expression early in life are thought to compensate for the immature adaptive immune system and to impact the dynamically developing skin microbiome.^24^

The role of hBDs in AD pathogenesis is unclear. While a positive impact may arise from their antimicrobial activities and improvement of skin permeability barrier at the tight junction level, hBDs can also act as immunomodulators by attracting and activating inflammatory cells, inducing secretion of inflammatory cytokines, and provoking itch, thus contributing to the pathogenesis of AD.^25^ Previous reports show higher levels of hBD-1 on skin surface samples from AD lesions compared to non-lesional skin.^26^ Here, we show that hBD-1 is upregulated on the skin of children who later developed AD compared to those who did not.

The S100A8/A9 heterodimer is expressed in epithelial cells in response to stress^27^ and can play both pro-inflammatory and anti-inflammatory roles (acting on neutrophils and suppressing dendritic cells, respectively),^27^ while also acting as an anti-microbial agent, called calprotectin.^28^ While S100A8 and S100A9 proteins are absent or minimally expressed in healthy epidermis, the synthesis of S100 proteins abruptly increases in epidermal keratinocytes in acute and chronic AD skin lesions.^29^ In acute AD, these proteins might contribute to chemotaxis of immune cells, particularly T-cells.^30^ Elevated serum levels of S100A8 reflect disease severity in canine AD.^31^ Moreover, S100A8/9 may be involved in inducing inflammation in AD triggered by IL-17A and house-dust mites.^32^ Finally, it has been reported that both lesional and non-lesional skin of AD patients are characterized by high S100A8/9 and low filaggrin expression, with further deficits in filaggrin expression in lesional skin.^33^

The presence of amino acids and derivatives on the skin surface are thought to be the result of proteolysis of filaggrin and related proteins in the SC.^34^ The results here confirm prior findings regarding a decrease in the amino acid concentration in the SC of AD patients compared to healthy individuals.^35^

Although non-lesional skin in AD patients is seemingly healthy skin as seen by macro-observations,^36^ it presents broad terminal differentiation defects and abnormalities in immune markers.^37^ Moreover, the non-lesional skin surface distinguishes AD with food allergy as a unique endotype and correlates with TEWL values.^38^ Here, both lesional and non-lesional skin on the face and the arm of AD patients had higher TEWL rates compared to corresponding skin sites of either the low- or high-risk non-AD group, but significance was only reached for the lesional skin sites. Raman measurements also revealed lower total amino acid content in AD patients (non-lesional skin) compared to the two non-AD groups. These observations indicate an intrinsic predisposition to barrier abnormalities and/or background immune activation in non-lesional AD patients. Further investigation into non-lesional sites is warranted and may prove useful as predictive indicators to late-onset eczema.

Finally, in this study machine learning models trained on the immune marker concentration differences on the arm between healthy/low-risk and AD-lesional skin could classify converters and non-converters. However, given the limited number of samples used to train (59 samples) and test (101 samples) the machine learning models, these models need to be validated on a larger dataset before one can be confident about the ability of these markers to predict future development of AD.

A potential limitation of this study is that information on AD development was reported by parents. Clinical examination and diagnosis by an expert physician would be more accurate. However, that would also add complexity in the study design since AD could develop at any time during the 12 months of follow-up.

These results show that in children 3-48 months of age, parameters such as the TEWL and SSH levels were not significantly different even between groups. Nevertheless, this study shows for the first time that in healthy children a combination of four skin surface biomarkers was associated with development of AD in the subsequent 12 months following measurements. Future research should focus on uncovering the mechanisms by which these biomarkers contribute to pathogenesis of AD.

