## Supplemental material for "Skin surface biomarkers are associated with future development of atopic dermatitis in children with family history of allergic disease"

### SUPPLEMENTAL METHODS

#### Statistical analysis

Outliers identified as falling in the interval $[q_{25} - 1.5 * IQR;\ q_{75} + 1.5 * IQR]$ were removed, where $q_{25}$ is the quantile at 25% per group, $q_{75}$ the quantile at 75% per group, and $IQR = q_{75}$ - $q_{25}$ the inter-quantile range per group.

To account for differences in sex, age, and birth method between groups, a linear statistical model of the calculated areas under the curve (AUCs) was built and adjusted with respect to these 3 variables according to:

$$X_{lm_{i,j}} \sim a_0 + a_1 * sex_i + a_2 * \ age_i + a_3 * \ birth\ method_i \qquad (1)$$

where $X_{lm_{i,j}}$ is variable *X (AUC, total amino acids, TEWL, or SSH measures)* as calculated by the linear model for subject *i* at measuring site *j* (arm or face), $sex_i$ is subject *i's sex*, $age_i$ is subject *I*'s age, $birth\ method_i$ is subject *i*'s birth mode of delivery, $a_k$ linear model's parameters,

and

$$X_{adjusted_{i,j}} \sim X_{lm_{i,j}} - \ a_0 - \ a_1 * \ sex_i - a_2 * \ age_i - a_3 * \ birth\ method_i \qquad (2)$$

where $X_{adjusted_{i,j}}$ is variable *X* adjusted with respect to sex, age, and birth method for subject *i* at measuring site *j* (arm or face). Group comparisons for the adjusted variables were tested using ANOVA and pairwise Bonferroni tests.

*Raman data analysis*

Raman spectra per risk group were plotted with respect to location in the stratum corneum (surface location is set to 0). Water profiles that did not reach a plateau in the viable epidermis were removed. Water resistance profiles were computed using water concentration profiles and corresponding TEWL values, as described by van Longtestijn et al.[19] To evaluate the difference between risk groups, we computed the AUC for each water resistance profile over the SC thickness (normalized depth between 0-1).

For the spectra collected in the fingerprint region, outliers of spectral stacks were determined using the keratin profiles and removed from subsequent analysis. The calculated concentration profiles were normalized by SC thickness for each subject. Total amino acid profiles were obtained from the sum of the amino acid concentration profiles (as above, calculated by SkinTools). Average total amino acid concentration profiles per group were computed by averaging data points at a certain depth for all subjects in a risk group. To evaluate the difference between the 3 risk groups, the AUC was computed for each total amino acid concentration profile in the top 80% of the SC, *i.e.* normalized depth between 0-0.8.

*Immune marker analysis*

Differential expression of immune markers was performed separately for the face and arm using a linear model that incorporated age, sex, and birth mode as covariates. Combined Z-score was generated by summing up the individual Z-scores of all measured markers. Correlation between different variables was performed using Spearman's rank correlation.

*Machine learning*

To gauge the ability of immune markers in predicting converters, 4 different machine learning models (Elastic Net, Random Forest, Support Vector Machine, XGBoost) were trained to predict AD-lesional subjects vs. healthy low-risk subjects using all marker data. Models underwent 3-fold cross-validation, repeated 10 times, for hyperparameter tuning. The trained models were then tested on predicting converters vs. non-converters.

**SUPPLEMENTAL TABLES**

**Table S1:** Location of AD lesional and healthy surrounding skin sites where swab samples were collected from in AD patients (Group C).

| Subject No. | Site 1 (Cheek) | Site 2 (Arm or non-cheek) | Site 3 (Surrounding) |
|---|---|---|---|
| 22 | Cheek (Lesion) | Antecubital fossa | Forehead |
| 24 | Cheek | Leg (Lesion) | Leg |
| 35 | Cheek (Lesion) | Antecubital fossa | Forehead |
| 56 | Cheek | Chest (Lesion) | Chest |
| 75 | Cheek | Hip (Lesion) | Abdomen |
| 76 | Cheek | Antecubital fossa (Lesion) | Antecubital fossa |
| 79 | Cheek | Cheek (Lesion) | Forehead |
| 82 | Cheek | Shoulder (Lesion) | Upper Arm |
| 86 | Cheek (Lesion) | Antecubital fossa | Forehead |
| 96 | Cheek | Collarbone (Lesion) | Collarbone |
| 99 | Cheek (Lesion) | Antecubital fossa | Forehead |
| 112 | Cheek (Lesion) | Antecubital fossa | Forehead |
| 120 | Cheek | Popliteal fossa (Lesion) | Popliteal fossa |
| 127 | Cheek (Lesion) | Antecubital fossa | Forehead |
| 132 | Cheek | Hand (Lesion) | Hand |
| 133 | Cheek | Temporal (Lesion, eye area) | Forehead |
| 134 | Cheek (Lesion) | Antecubital fossa | Forehead (healthy surrounding) |
| 136 | Cheek (Lesion) | Antecubital fossa | Forehead |
| 137 | Cheek (Lesion) | Antecubital fossa | Forehead |
| 139 | Cheek (Lesion) | Antecubital fossa | Forehead |
| 144 | Cheek | Antecubital fossa (Lesion) | Antecubital fossa |
| 150 | Cheek | Volar forearm (Lesion) | Volar Forearm |
| 151 | Cheek | Popliteal fossa (Lesion) | Popliteal fossa |
| 152 | Cheek | Popliteal fossa (Lesion) | Popliteal fossa |
| 155 | Cheek | Popliteal fossa (Lesion) | Popliteal fossa |
| 156 | Cheek | Antecubital fossa (Lesion) | Antecubital fossa |
| 157 | Cheek (Lesion) | Antecubital fossa | Forehead |
| 158 | Cheek (Lesion) | Antecubital fossa | Forehead |
| 159 | Cheek (Lesion) | Antecubital fossa | Forehead |
| 160 | Cheek | Volar forearm (Lesion) | Volar forearm |

**Table S2:** Statistical comparisons (*p* values) of the biophysical and immune marker concentration parameters between converters and non-converters in the high-risk group for the two sites measured using a linear model. Group B: healthy/high-risk children with family history of allergic disease; converters: children in Group B who developed AD within 12 months after skin assessment; non-converters: children in Group B who did not develop AD within 12 months after skin assessment

| | **Arm** | **Face** |
|---|---|---|
| **TEWL** | 0.961 | 1.000 |
| **Skin surface hydration** | 0.967 | 1.000 |
| **Immune marker Z-score** | **0.036** | 0.827 |
| **CCL17** | NA | 1.000 |
| **CCL2** | 0.868 | 1.000 |
| **CCL22** | 1.000 | 1.000 |
| **CCL27** | 1.000 | 1.000 |
| **CXCL1/2** | NA | 1.000 |
| **hBD-1** | 0.430 | 0.250 |
| **IL-18** | 0.994 | 0.997 |
| **IL-1α** | 0.471 | 0.780 |
| **IL-1β** | 0.549 | 0.999 |
| **IL-1RA** | 0.113 | 1.000 |
| **IL-36γ** | 0.381 | 0.966 |
| **IL-8** | 0.938 | 1.000 |
| **S100A8/9** | 0.314 | 0.986 |
| **VEGF-A** | 1.000 | 0.215 |

AD, atopic dermatitis; TEWL, transepidermal water loss; CCL, chemokine (C-C motif) ligand; CXCL, chemokine (C-X-C motif) ligand; hBD-1, human β-defensin-1; IL, interleukin; IL-1RA, IL-1 receptor antagonist; VEGF-A, vascular endothelial growth factor A.

**Table S3:** Combined Z-scores were generated for the concentrations of immune markers measured on the arm with different combination of markers sorted on p values indicated in Table S1. These combined Z-scores were compared between converters and non-converters using a linear model. A minimum of 4 immune markers (IL-1RA, S100A8/9, IL-36γ, and hBD-1) was required to reach statistical significance ($p<0.05$) separating converters and non-converters. Group B: healthy/high-risk children with family history of allergic disease; converters: children in Group B who developed AD within 12 months after skin assessment; non-converter, ns: children in Group B who did not develop AD within 12 months after skin assessment.

| Markers, n | *p* value |
|---|---|
| 1 | 0.113 |
| 2 | 0.051 |
| 3 | 0.054 |
| 4 | **0.036** |
| 5 | **0.027** |
| 6 | **0.008** |
| 7 | **0.006** |
| 8 | **0.007** |
| 9 | **0.011** |
| 10 | **0.012** |
| 11 | **0.018** |
| 12 | **0.033** |
| 13 | **0.033** |
| 14 | **0.034** |

AD, atopic dermatitis; hBD-1, human β-defensin-1; IL, interleukin; IL-1RA, IL-1 receptor antagonist.

**Table S4:** Odds ratios and comparisons of the biophysical and immune marker parameters between converters and non-converters. Group B: healthy/high-risk children with family history of allergic disease; converters: children in Group B who developed AD within 12 months after skin assessment; non-converters: children in Group B who did not develop AD within 12 months after skin assessment.

**Arm**

| | **Odds Ratio** | **2.5% CI** | **97.5% CI** | ***p* value** |
|---|---|---|---|---|
| Immune marker Z-score | 19.216 | 3.510 | 186.062 | **0.002** |
| Age | 0.427 | 0.067 | 2.464 | 0.338 |
| Sex | 3.061 | 0.550 | 25.285 | 0.230 |
| Birth mode | 1.160 | 0.217 | 7.161 | 0.864 |
| Skin surface hydration | 2.790 | 0.461 | 28.138 | 0.309 |
| TEWL | 1.225 | 0.161 | 13.057 | 0.850 |

**Face**

| | **Odds Ratio** | **2.5% CI** | **97.5% CI** | ***p* value** |
|---|---|---|---|---|
| Immune marker Z-Score | 9.097 | 1.532 | 67.540 | **0.018** |
| Age | 0.345 | 0.048 | 2.151 | 0.259 |
| Sex | 2.310 | 0.466 | 13.138 | 0.310 |
| Birth mode | 0.814 | 0.141 | 5.184 | 0.817 |
| Skin surface hydration | 0.985 | 0.184 | 6.425 | 0.986 |
| TEWL | 1.340 | 0.209 | 10.079 | 0.760 |

AD, atopic dermatitis; CI, confidence interval; TEWL, transepidermal water loss.

**Table S5:** Linear model used to compare biomarkers between groups B and C to group A on the arm and the face. Group A: Healthy/low-risk, Group B: Healthy/high-risk, and Group C: AD patients. AD, atopic dermatitis; CCL, chemokine (C-C motif) ligand; CXCL, chemokine (C-X-C motif) ligand; hBD-1, human β-defensin-1; IL, interleukin; IL-1RA, IL-1 receptor antagonist; NA, not available; SSH, skin surface hydration; TEWL, transepidermal water loss; VEGF-A, vascular endothelial growth factor A.

| **Arm** | **Group C (Lesional)** | **Group C (Non-lesional)** | **Group C (Healthy surrounding)** | **Group B** |
|---|---|---|---|---|
| **TEWL** | **<0.001** | 0.179 | 0.934 | 1.000 |
| **SSH** | **<0.001** | 1.000 | 0.928 | 0.964 |
| **Immune marker Z-score** | **<0.001** | 1.000 | 0.905 | 1.000 |
| **CCL-17** | NA | NA | NA | NA |
| **CCL-2** | 0.575 | 0.801 | 0.993 | 0.977 |
| **CCL-22** | **<0.001** | 0.998 | 0.754 | 1.000 |
| **CCL-27** | **<0.001** | 1.000 | 1.000 | 1.000 |
| **CXCL1/2** | NA | NA | NA | NA |
| **hBD-1** | 0.898 | 0.970 | 1.000 | 0.769 |
| **IL-18** | 0.053 | 0.857 | 0.928 | 0.998 |
| **IL-1α** | 0.120 | 0.928 | 0.294 | 1.000 |
| **IL-1β** | 1.000 | 0.999 | 1.000 | 0.783 |
| **IL-1RA** | **<0.001** | 0.485 | 0.991 | 1.000 |
| **IL-36γ** | **<0.001** | 0.629 | 0.929 | 0.973 |
| **IL-8** | **<0.001** | 0.986 | 0.995 | 0.994 |
| **S100A8/9** | **<0.001** | 0.999 | 0.850 | 0.718 |
| **VEGF-A** | 1.000 | 0.912 | 0.925 | 0.997 |

| **Face** | **Group C (Lesional)** | **Group C (Non-lesional)** | **Group C (Healthy surrounding)** | **Group B** |
|---|---|---|---|---|
| **TEWL** | 0.225 | 0.999 | 1.000 | 0.983 |
| **SSH** | 0.166 | 0.847 | **0.019** | 0.704 |
| **Immune marker Z-score** | **<0.001** | 0.928 | 1.000 | 0.926 |
| **CCL-17** | **<0.001** | 1.000 | 0.998 | 0.998 |
| **CCL-2** | **0.035** | 1.000 | 0.971 | 1.000 |
| **CCL-22** | 0.660 | 1.000 | 0.110 | 1.000 |
| **CCL-27** | **<0.001** | 1.000 | 0.944 | 1.000 |
| **CXCL1/2** | **<0.001** | 0.159 | 0.993 | 0.997 |
| **hBD-1** | 0.999 | 0.187 | **<0.001** | 0.192 |

| | | | | |
|---|---|---|---|---|
| **IL-18** | **0.003** | 1.000 | 1.000 | 1.000 |
| **IL-1α** | 0.994 | 0.186 | 0.944 | 0.967 |
| **IL-1β** | 0.737 | 0.291 | 0.366 | 0.500 |
| **IL-1RA** | 0.930 | 0.385 | 0.051 | 0.997 |
| **IL-36γ** | 0.564 | 0.875 | 0.678 | 0.985 |
| **IL-8** | **<0.001** | 0.925 | 0.541 | 1.000 |
| **S100A8/9** | **0.010** | 0.975 | 0.993 | 0.773 |
| **VEGF-A** | 0.962 | 0.592 | 0.129 | 1.000 |

### SUPPLEMENTAL FIGURES

**Figure S1: Average profile of resistance to water transport in the SC for (a) the groups tested (Group A: Healthy/low-risk, Group B: Healthy/high-risk, and Group C: AD patients, non-lesional site), and (b) AD converters vs. non-converters in the high-risk group.** Error bars indicate one standard deviation. The profile depth is normalized to the SC thickness as calculated from the water concentration profiles. Pairwise t-test comparisons show significant differences in the area under the curve between groups A, B, and C (adjusted p-value<0.5), but no significant difference between converters and non-converters. AD, atopic dermatitis; SC, stratum corneum.

**a** **b**

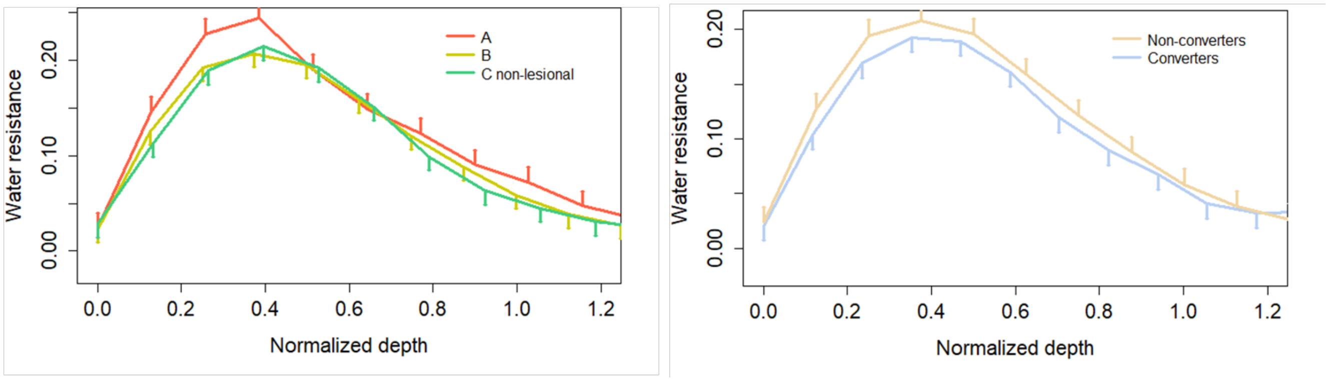

**Figure S2: Average concentration profile of total amino acids in the SC for (a) the groups tested (Group A: Healthy/low-risk, Group B: Healthy/high-risk, and Group C: AD patients, non-lesional site), and (b) AD converters vs. non-converters in the high-risk group.** Error bars indicate one standard deviation. The profile depth is normalized to the SC thickness as calculated from the water concentration profiles. Pairwise T-test comparison show significant differences in the area under the curve between groups A and C (adjusted p-value<0.05), but no significant difference between groups A and B, nor between groups B and C or between converters and non-converters in group B. AD, atopic dermatitis; SC, stratum corneum.

**a**

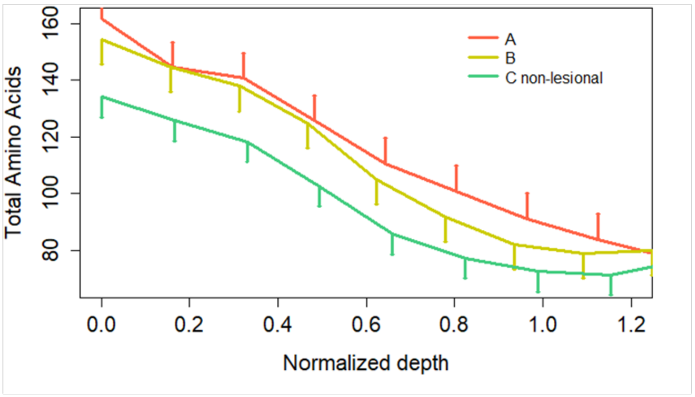


**b**

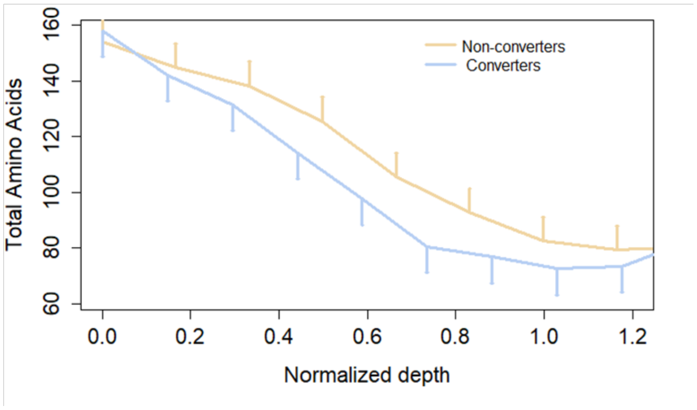

**Figure S3.** Typical appearance of skin surface on the arm site for the (a) Group A: heathy/low-risk group, (b) Group B: healthy/high-risk group, (c) Group C: non-lesional skin site in the AD group, and (d) lesional skin in the AD group. AD, atopic dermatitis.

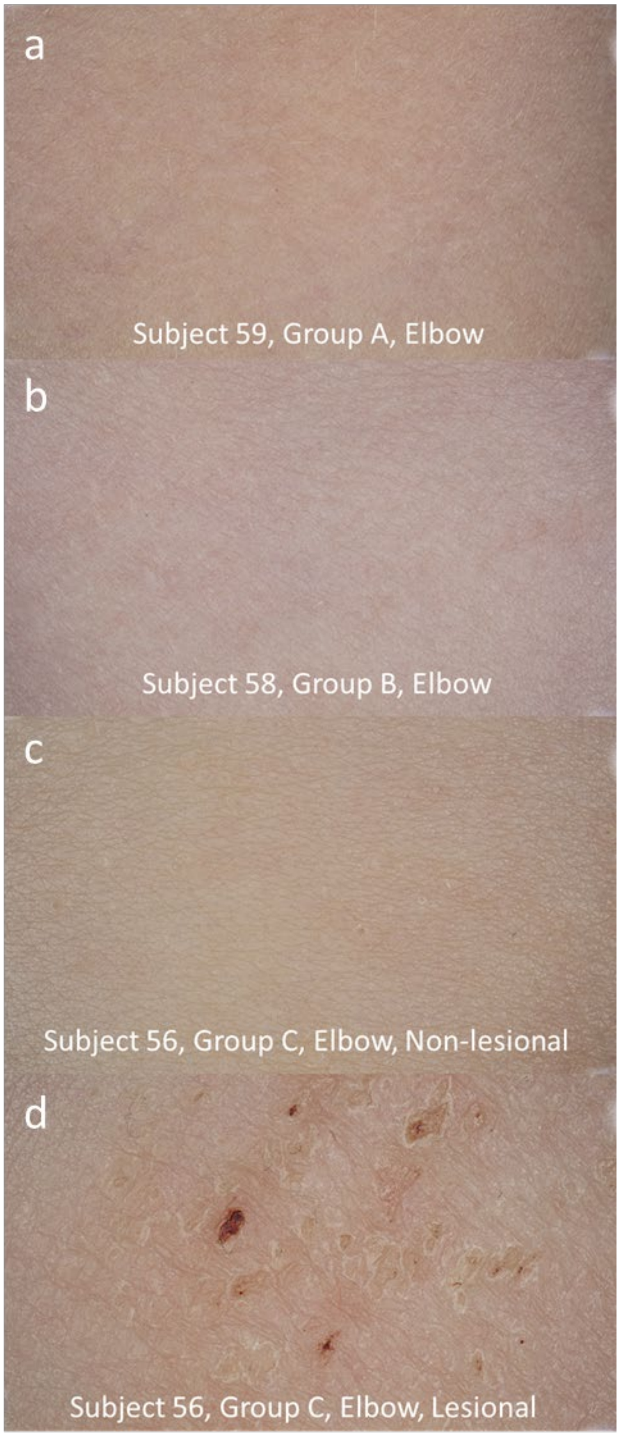

**Figure S4: Correlation between TEWL and skin surface marker concentrations.** Spearman's rank correlation coefficients (indicated by the bar on the right) were generated between TEWL and the skin surface marker concentration on the (a) face and (b) arm site in healthy/low-risk (A), healthy/high-risk (B), AD non-lesional (C), and AD lesional site (C lesion). AD, atopic dermatitis; CCL, chemokine (C-C motif) ligand; CXCL, chemokine (C-X-C motif) ligand; hBD-1, human β-defensin-1; IL, interleukin; IL-1RA, interleukin-1 receptor antagonist; SSH, skin surface hydration; TEWL, transepidermal water loss; VEGF-A, vascular endothelial growth factor A. Question mark signs indicate too few values to perform statistics (most samples measured showed values below the detection limit of the assay).

**a**

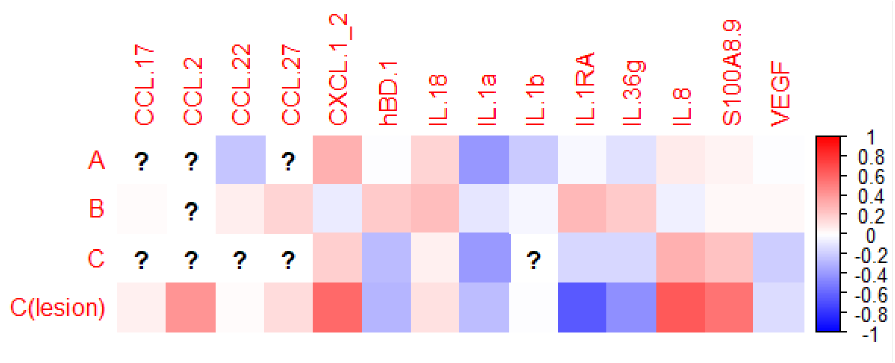


**b**

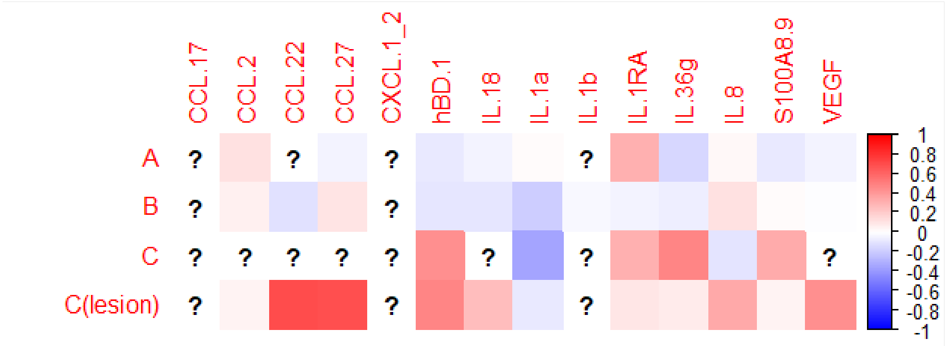

**Figure S5: Skin surface concentration of hBD1 decreases with age after birth.** Spearman correlation coefficients are -0.26 (p<00.1) for the arm and -0.31 (p<0.01) for the face sites. hBD-1, human β-defensin-1. Group A: healthy/low-risk children with no family history of allergic disease; Group B: healthy/high-risk children with family history of allergic disease; Group C: children with active AD lesions (local SCORAD index of 4-8).

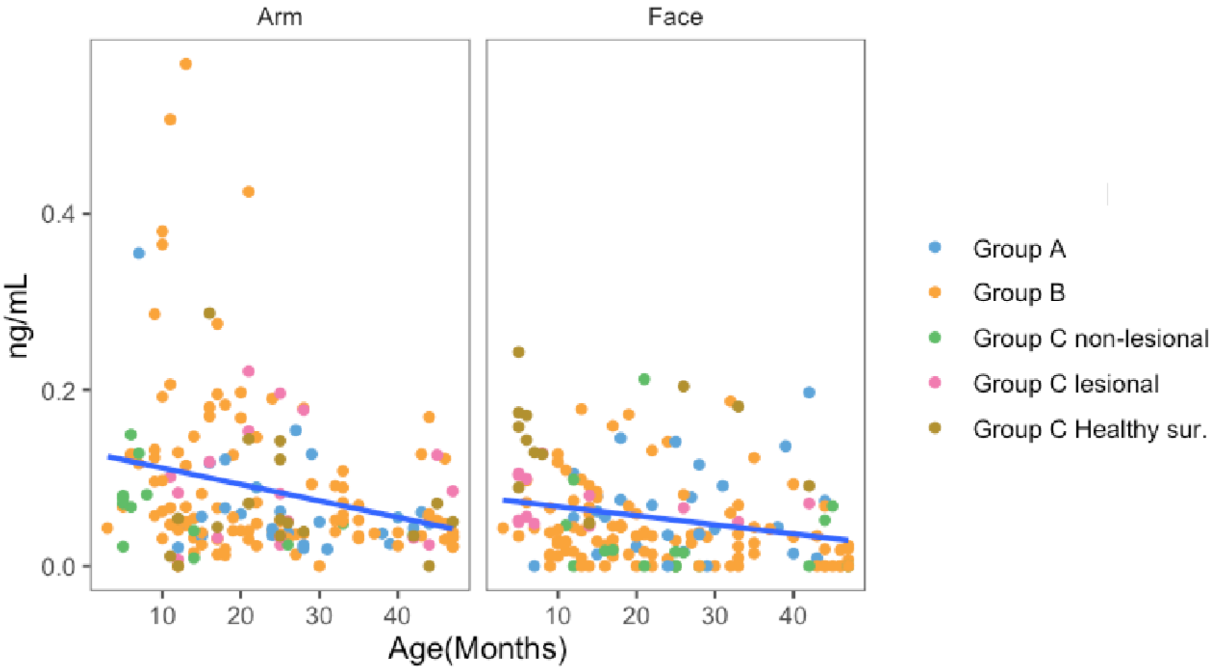
