## Supplementary material for "Skin surface biomarkers are associated with future development of atopic dermatitis in children with family history of allergic disease": Table 1

**TABLES**

**Table 1: Demographic and clinical data.**

Group A: healthy/low-risk children with no family history of allergic disease; Group B: healthy/high-risk children with family history of allergic disease; Group C: children with active AD lesions (local SCORAD index of 4-8); converters: children in Group B who developed AD within 12 months after skin assessment; non-converters: children in Group B who did not develop AD within 12 months after skin assessment.

| | Group A (n=29) | Group B (n=101) | Group C (n=30) | Total (n=160) | Group B converters (n=11) | Group B non-converters (n=71) |
|---|---|---|---|---|---|---|
| **Sex** | | | | | | |
| female | 16 (55.2%) | 57 (56.4%) | 11 (36.7%) | 84 (52.5%) | 6 (54.5%) | 51 (56.7%) |
| male | 13 (44.8%) | 44 (43.6%) | 19 (63.3%) | 76 (47.5%) | 5 (45.5%) | 39 (43.3%) |
| **Mode of Birth** | | | | | | |
| C-section | 10 (34.5%) | 29 (28.7%) | 8 (26.7%) | 47 (29.4%) | 3 (27.3%) | 26 (28.9%) |
| Vaginal | 19 (65.5%) | 72 (71.3%) | 22 (73.3%) | 113 (70.6%) | 8 (72.7%) | 64 (71.1%) |
| **Age (months)** | | | | | | |
| Mean (SD) | 25.9 (10.7) | 23.1 (12.6) | 20.6 (13.4) | 23.1 (12.5) | 23.5 (13.9) | 23.0 (12.6) |
| Median | 25 | 20 | 19 | 21 | 18 | 20 |
| [Min, Max] | [7, 44] | [3, 47] | [5, 47] | [3, 47] | [6, 44] | [3, 47] |
| **TEWL ($gm^{-2}h^{-1}$)** | | | | | | |
| Mean (SD) | | | | | | |
| Arm | 13.7 (6.2) | 14.0 (7.0) | 18.3 (7.6)* | | 11.8 (2.6) | 14.7 (8.0) |
| Face | 17.2 (10.1) | 17.1 (6.1) | 19.0 (7.0)* | | 16.4 (5.7) | 17.0 (6.2) |
| **SSH (AU)** | | | | | | |
| Mean (SD) | | | | | | |
| Arm | 38.4 (8.7) | 40.9 (13.4) | 38.6 (13.1)* | | 38.9 (12.5) | 42.6 (14.1) |
| Face | 32.9 (9.7) | 36.4 (11.8) | 35.8 (16.3)* | | 37.8 (6.4) | 38.1 (12.4) |

AD, Atopic Dermatitis; SCORAD, SCORing Atopic Dermatitis; SD, standard deviation; TEWL, Transepidermal water loss; AU, arbitrary units.

* Measurements on non-lesional skin site.
